# Female Túngara Frogs Discriminate against the Call of Males Infected by Chytridiomycosis

**DOI:** 10.1101/2024.08.06.606873

**Authors:** Sofía Rodríguez-Brenes, Sylvia F. Garza, Michael J. Ryan

## Abstract

Species worldwide are disappearing in the most devastating mass extinction in human history and one of the six most profound extinctions in the history of life. Amphibians are greatly affected, approximately one third of living species are threatened, and many others are extinct. One of the main causes of amphibian species extinctions and population declines is the emerging infectious disease chytridiomycosis, caused by the fungus *Batrachochytrium dendrobatidis* (Bd). Although some species are somewhat tolerant of the disease, the non-lethal effects of the infection with Bd and their short or long term consequences are poorly understood. In these species there is the potential for behavioral responses to mitigate the spread of the fungus. Here we show that in túngara frogs, infection status influences the males’ mating calls. These infection-induced changes in the quality of males’ mating calls ultimately reduce the calls’ attractiveness to females making females less likely to respond to and thus mate with infected males. More broadly, our results imply that females might avoid mating with disease-infected males by assessing the acoustic signal only, and that such recruitment of behavioral responses might potentially ameliorate some of the effects of this sixth mass extinction.

**Lay summary:** Chytridiomycosis is an amphibian disease well known for its lethal effects. Túngara frogs are infected in nature, but seem to be resistant to the disease. Here we show that chytridiomycosis has non-lethal behavioral effects on túngara frogs. Females discriminate against infected males by assessing only their acoustic signal. The mating call of a male that is not infected with the disease is more attractive to females than the call of that same male when he is infected.

## Introduction

Amphibian species are declining in the largest mass extinction in human history and one of the six most profound extinctions in the history of life (Wake and Vredenburg 2008; Collins and Crump 2009; Kolbert 2014; Ceballos et al. 2015). An emergent disease, chytridiomycosis, has had devastating effects on amphibian populations across the world and is one of the main contributors to this catastrophic loss of biodiversity. Chytridiomycosis, caused by the chytrid fungus, *Batrachochytrium dendrobatidis* (Bd), is well known for its lethal effects. Bd disrupts osmoregulatory functions of amphibian skin ultimately causing cardiac arrest (Voyles et al. 2009), and it also inhibits the host’s immune system (Fites et al. 2013). Amphibian hosts vary in their defense mechanisms to fight the infection from Bd; their skin microbiota may function as a first barrier against the fungi (Harris et al. 2009), and immune responses specific to Bd have also evolved in some amphibian populations (Savage and Zamudio 2011; Bataille et al. 2015; Ellison et al. 2015; Kosch et al. 2016). In nature, some species, including the túngara frog (*Physalaemus* [*Engystomops*] *pustulosus*), are infected with Bd but are thriving and do not show any of the lethal symptoms of the disease (Rodríguez-Brenes et al. 2016). In a few species behavioral effects due to the infection with Bd have been observed at the larval and adult stage. In two species of frogs, *Hyla chrysoscelis,* and *Anaxyrus fowleri,* tadpoles’ foraging behavior decreased when infected with Bd (Venesky et al. 2009). In adults of *H. japonica*, changes in the characteristics of male’s mating call have been associated with the infection, although fitness consequences for these males were not investigated (An and Waldman 2016). Other physiological effects in adults of *Lithobates pipiens* infected with Bd in the lab, suggest an increase in reproductive effort induced by this pathogen infection (Chatfield et al. 2013). The fitness consequences of these non-lethal effects also are unknown.

Amphibians can contract Bd through contact with fungal spores in the environment and from contact with other infected individuals (Rachowicz and Vredenburg 2004). Bd can be transmitted, for example, in both directions between tadpoles and adults and between conspecifics and heterospecifics (Fernández-Beaskoetxea et al. 2016). Given that Bd infects the skin, adult-adult contact is also assumed to mediate direct transmission of the disease. An especially dangerous venue for Bd transmission should occur during sex. In most frog species females assess the male’s acoustic sexual signal and decide with whom to mate. Male túngara frogs produce a whine, which females use to recognize conspecifics, and then add one or more chucks, which further increases their attractiveness to females (Rand and Ryan 1981). Mate choices are usually followed by a ‘love hug’ or amplexus, which often lasts for hours allowing sufficient time for the transmission of the pathogen. Frog calls in general, and túngara calls in particular, are energetically very costly (Ryan 1985). Thus a male’s immune system allocating resources and energy to cope with a pathogen infection could influence the amount of energy devoted to calling. This has been shown in other amphibians: in the treefrog, *Hypsiboas prasinus*, the presence of helminth parasites decreased the calling rate of males (Madelaire et al. 2013), and infection with Bd, in the frog *Litoria rheocola,* decreases the probability of calling in males with low body condition (Roznik et al. 2015). In neither of these cases were the effects of these behavioral changes tested, e.g. if females can detect the changes in the acoustic signal due to the infection with the disease, and if this effect is sufficient to alter the attractiveness the males’ call.

Here we asked if female túngara frogs are able to discriminate between infected and uninfected males by attending to the frog’s acoustic signal only, i.e., the mating call isolated from other aspects of the male’s phenotype. We hypothesized that infection by Bd should influence components of the males’ sexual mating signal. To test this, we recorded the mating calls of the same males in infected and uninfected states, and presented the calls to females in phonotaxis tests. This approach is superior to comparing the call of different individuals that are either infected or uninfected as it controls for other variables of the males’ genotype and phenotype that affect the production of sound.

## Methods

### Bd sampling

To test for infection with Bd we swabbed the frog’s ventral area with a dry swab (Medical Wire and Equipment, model MW110) following established protocols (Hyatt et al. 2007). Swabs were analyzed by qPCR following Boyle et al. (2004) and modified by Kriger et al. (2006). The swabs were stored in 70% ethanol and analyzed by real-time quantitative PCR (qPCR) using TaqMan® Fast Advanced, Master Mix (Applied Biosystems) and a StepOnePlus^TM^ system, with a dilution series of genomic DNA from strain JEL423 as our standard reference.

### Male recordings

We collected males in Gamboa, Panama in June 2014 and transported them to the University of Texas at Austin, USA. We kept them in captivity in our túngara frog colony facilities. To test for infection with Bd, we swabbed each male prior to recording their mating call. We recorded the males in a sound attenuation chamber under controlled conditions. Each male was placed in a small container with mesh on the sides to allow sound propagation, and with a bowl of water from where he could call. The microphone (Sennheiser ME66) was 50 cm from the bowl of water. We recorded the males’ call with a Zoom H6 recorder at a sampling rate of 44.1 kHz and 16 bit per sample. To record the amplitude of the call, at the beginning of each file we recorded a tone of 780 Hz, which approximates the dominant frequency of the whine, at an amplitude of 82 dB SPL (re. 20 µPa) at the same position of the bowl of water (50 cm) from where the males call. To stimulate the males to call, we broadcast bouts of a chorus of túngara males recorded in the wild, with silence intervals to be able to record the males’ call.

The three males used in the experiments were infected when collected, thus they were also infected prior to the first recording. To clear the males of Bd, we treated them with a solution of 0.0025% Itraconazole once a day for 5 min, for 6 consecutive days (Brannelly 2014). After each 5 min treatment we placed them in a clean container to avoid reinfection. Males were swabbed before the second recording to confirm they were cleared of the infection. We recorded them under the same conditions and using the same procedure described above. In this way we were able to control for variation in the call parameters due to environmental factors, and also aspects of the call related with the males’ phenotype.

To capture the maximum amplitude at which the males were able to call, we chose four calls with the highest amplitude of each recording to measure their spectral and temporal properties. From each male we chose calls of both recordings, infected and uninfected with Bd, with the same structure (whine vs whine, or whine-chuck vs whine-chuck). The parameters we measured are represented in Figure 1: the time to half rise (duration from the onset of the call to one-half the maximum amplitude during the rise), the time to half fall (time from the beginning of the call to the point at one-half the maximum amplitude during the fall), rise time (time from the whine’s onset to its maximum amplitude), fall time (time from the whine’s maximum amplitude to the end of the whine), time to half frequency (whine’s duration from the onset to its mid-frequency), final frequency (whine’s frequency at the end of the whine), maximum frequency (maximum frequency of the whine’s fundamental), dominant frequency (dominant frequency of the call), and amplitude (root-mean-square amplitude). For sound analysis we used Raven Pro 1.4 (Bioacoustics Research Program, Cornell Lab of Ornithology, Ithaca, NY). We performed a principal component analysis (PCA) in R (R Core Team 2015) to visualize differences in the call parameters as a function of Bd infection status.

**Fig. 1.**
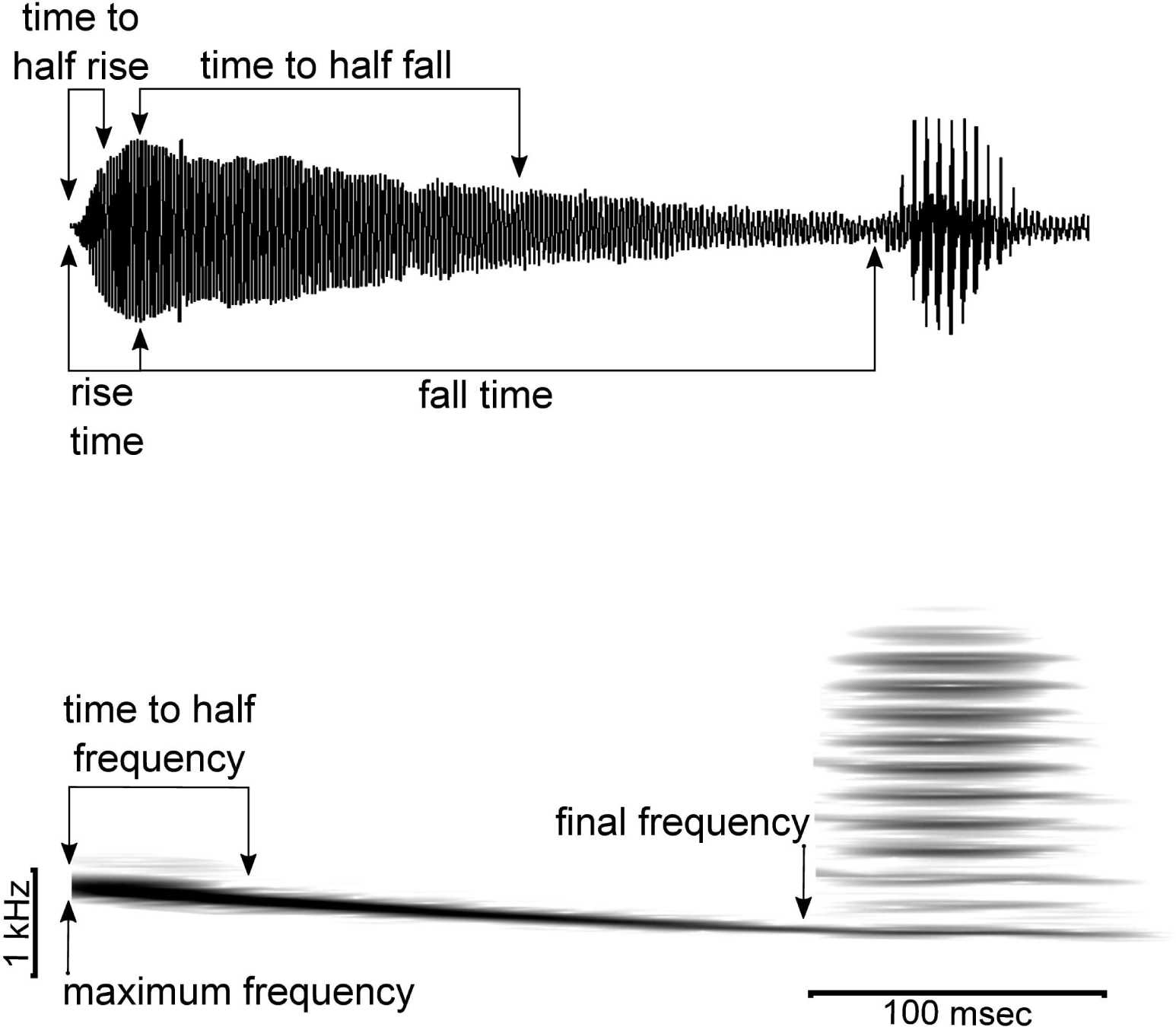
Túngara frog males’ mating call. Sonogram and spectrogram of a whine and a chuck, and the call variables measured for this study.

### Female phonotaxis test

To determine if females can discriminate against the calls of males when they are infected with Bd, we tested females in a standard two-choice phonotaxis test (Ryan and Rand 1990). We performed these tests in Gamboa, Panamá at the Smithsonian Tropical Research Institute (STRI) from August to December of 2015. We collected pairs in amplexus and we tested the females that same night. We placed females in the center of a sound attenuation chamber (dimensions: 2.7 x 1.8 m), equidistant from a pair of speakers positioned at 180° at opposite ends of the chamber. We video recorded the tests using an infrared camera placed on the ceiling of the sound chamber. To avoid recapture of females, we toe-clipped them at the end of the night (Guimarães et al. 2014). We returned the pairs the next night to where we had collected them.

For the phonotaxis tests we used the same four calls for each of the three males that we analyzed and described above. We will refer to the three males as A, B, and C. For the test stimuli, we looped these four calls per recording from each of the three males. For an accurate playback of the absolute amplitude at which the males called during the recording, we calibrated each speaker using the tone recorded at the beginning of each file. The tone of each file was played back and measured at 82 dB SPL (re. 20 µPa) at the center of the acoustic chamber. The pair of stimuli was broadcast at a rate of one call per 2 s, alternating from each speaker. We allowed the female to acclimatize for 2 min under an inverted mesh funnel. The female was then released, and her preferred stimulus was recorded. We scored the female’s preference when the female advanced to within 10 cm in front of one of the speakers. Testers were blind to the treatments broadcast from each speaker.

We tested three versions of the call of the same three males, ‘natural’, ‘normalized’, and ‘inverted’ amplitude calls. The natural calls were playback at the original absolute amplitude at which they were recorded. The normalized and inverted calls, were modified using Adobe Audition 3 (Copyright © 2007 Adobe Systems Incorporated). For the normalized calls, we modified each pair so that they would have the same peak amplitude. For the inverted calls, we inverted the difference of peak amplitude of the original natural calls of each male (Fig 1). The order of the males’ calls was randomized every day. In August 2015, we tested 67 females with the natural calls. All the females completed all three tests, corresponding with the pair of calls from the three males. From October to December 2015, we tested a separate set of 87 females with the normalized and inverted calls, therefore each female completed six tests.

To determine whether the Bd status of males had an effect on female preference for their calls we used a general linear mixed effects models (glmer) with a binomial family and logit link function in R (R Core Team 2015) using the lme4 package (Bates et al. 2015). Models included male and female identity as a random factor, and the size (snout-vent length), mass, and Bd status of females as fixed factors. We expected the variation in males’ mating call due to infection with Bd to have a small effect overall on mate choice preference in females. To detect a strength of preference of 0.7 with a statistical power of 0.8 we required a sample size of 42, and of 56 individuals to increase the statistical power to 0.9. Female phonotaxis tests are relatively non-invasive, and the high abundance of túngara frogs in our field sites allow us to have sample sizes (naturals calls, n=67; normalized and inverted calls, n=87) to obtain results with a high statistical power.

## Results

### Bd infection effects on males’ acoustic signals

The mating calls of males when infected with Bd had a lower amplitude and higher dominant frequency than the calls of the same males when not infected (Table 1). When we plotted the multivariate properties of the six calls in a principal component analysis, the first principal component (PC1) explained 38% of the variation amongst the calls (Fig. 2, Table 2). Along this axis calls tended to group by infection status rather than individual. In addition, amplitude and dominant frequency of the whine had the higher loads on PC1. We then asked if these differences in the characteristics of the call affect the males’ call attractiveness to females.

**Fig 2.**
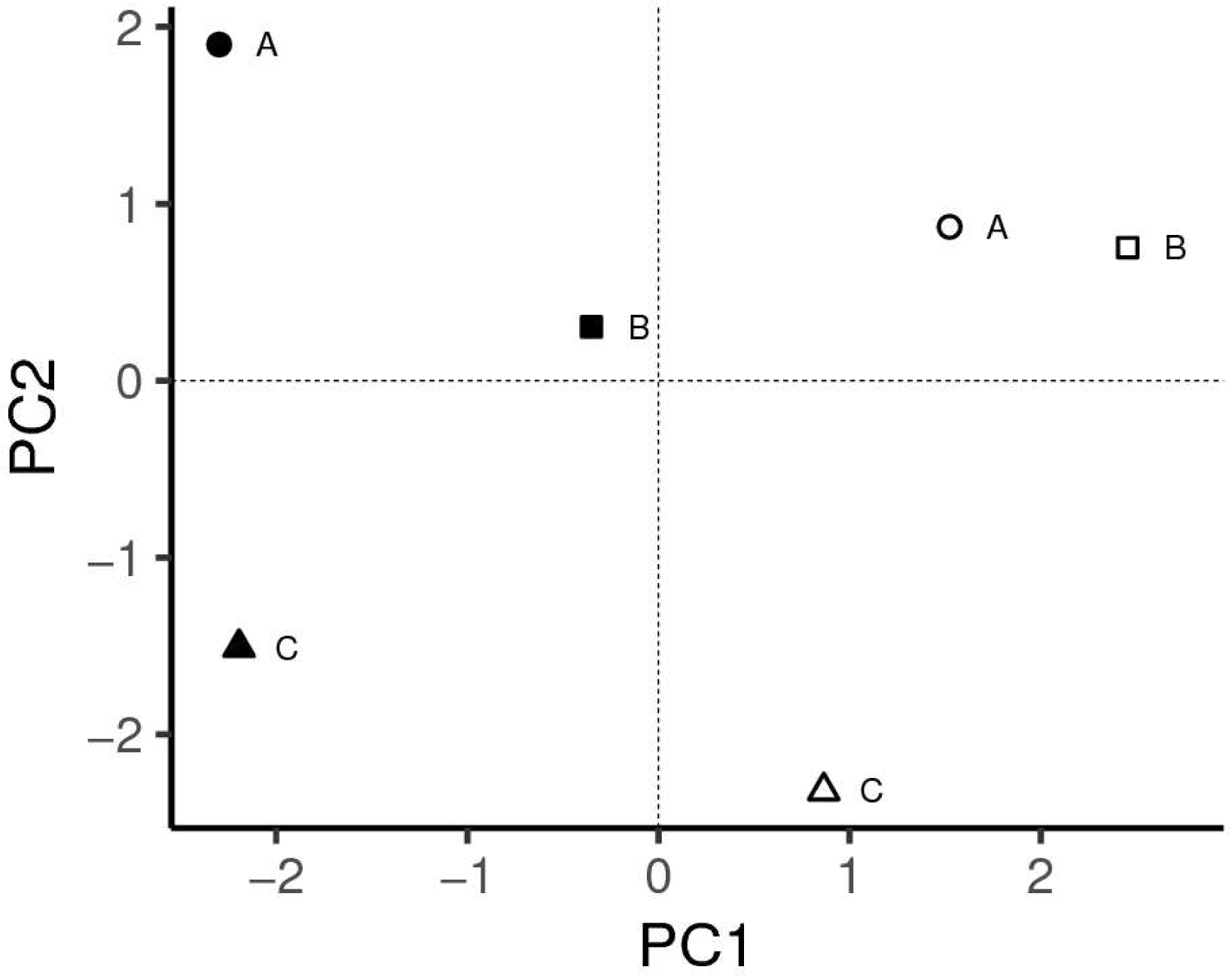
Principal components analysis of the multivariate properties of mating calls of túngara frogs. Call properties of mating calls of three males: A, B and C, in their infected with Bd (filled symbols) vs uninfected with Bd (open symbols) status. First principal component (PC1) had high loadings for amplitude of the call and the whine’s dominant frequency.

**Table 1.** Average values for measured properties of the mating calls of three túngara males when infected with Bd (+) vs when uninfected with Bd (-).

| Male | Bd infection status | Time to half frequency (s) | Dominant frequency (Hz) | Initial frequency (Hz) | Final frequency (Hz) | Time to half fall (s) | Time to half rise (s) | Rise time (s) | Fall time (s) | Amplitude |
| --- | --- | --- | --- | --- | --- | --- | --- | --- | --- | --- |
| A | + | 0.137 | 861.3 | 882.8 | 430.7 | 0.042 | 0.008 | 0.025 | 0.272 | 813.20 |
| A | - | 0.069 | 753.6 | 818.2 | 430.7 | 0.100 | 0.006 | 0.021 | 0.341 | 1774.10 |
| B | + | 0.064 | 947.5 | 1098.2 | 516.8 | 0.028 | 0.005 | 0.014 | 0.281 | 838.38 |
| B | - | 0.048 | 775.2 | 861.3 | 473.7 | 0.028 | 0.005 | 0.014 | 0.281 | 938.25 |
| C | + | 0.078 | 861.3 | 947.5 | 516.8 | 0.128 | 0.003 | 0.038 | 0.248 | 873.43 |
| C | - | 0.050 | 818.2 | 861.3 | 559.8 | 0.193 | 0.004 | 0.011 | 0.292 | 1806.20 |

**Table 2.** Principal component analysis (PCA) summary of the multivariate properties of the mating calls of túngara males measured in this study. Amplitude and dominant frequency of the whine have the highest load in the first principal component.

|  | PC1 | PC2 | PC3 | PC4 | PC5 |
| --- | --- | --- | --- | --- | --- |
| Standard deviation | 0.966 | 1.594 | 1.471 | 0.934 | 0.749 |
| Proportion of variance | 0.386 | 0.254 | 0.216 | 0.087 | 0.056 |
| Cumulative proportion | 0.386 | 0.640 | 0.857 | 0.944 | 1.000 |
| **Loading values** |  |  |  |  |  |
| Time to half frequency | 0.234 | 0.134 | -0.444 | 0.349 | 0.611 |
| Dominant frequency | **0.432** | -0.113 | 0.267 | 0.251 | 0.262 |
| Initial frequency | 0.405 | -0.065 | 0.383 | 0.157 | -0.168 |
| Final frequency | -0.013 | -0.477 | 0.436 | -0.060 | 0.123 |
| Time to half fall | -0.313 | -0.394 | 0.119 | 0.418 | 0.275 |
| Time to half rise | 0.296 | 0.454 | 0.210 | -0.070 | 0.264 |
| Rise time | 0.205 | -0.209 | -0.388 | 0.545 | -0.503 |
| Fall time | -0.386 | 0.389 | 0.050 | 0.196 | -0.043 |
| Amplitude | **-0.466** | 0.041 | 0.171 | 0.279 | 0.209 |

### Females discriminate against Bd infected males

In the first experiment using the natural calls, 67 females responded to three tests, one for each pair of calls from the three males, for a total of 198 phonotaxis choices. In this experiment, females discriminated between Bd-infected and uninfected males (Fig. 3A, Table 3) preferring the calls of the males when non-infected.

**Fig 3.**
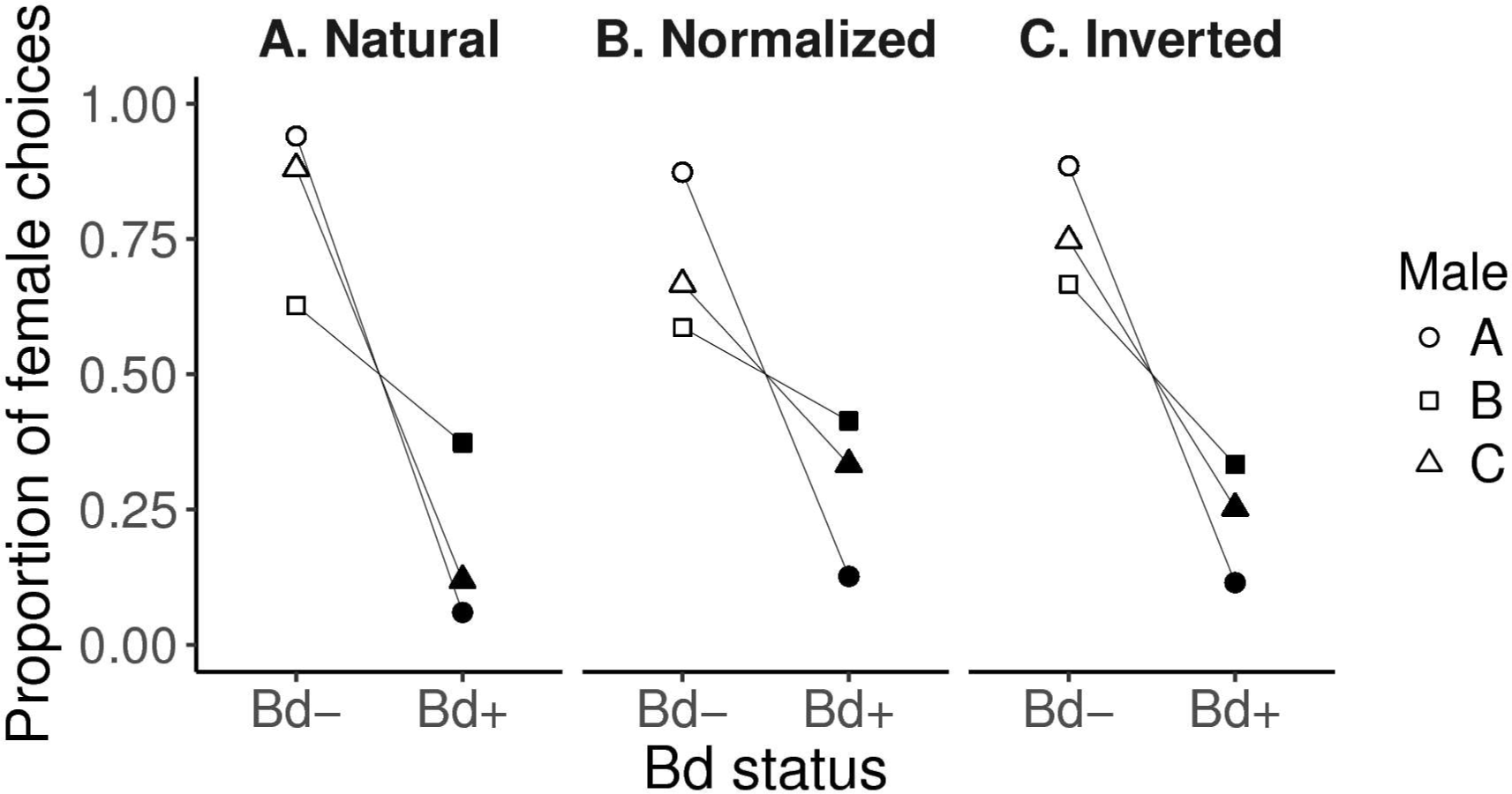
Female preference for calls of males uninfected and infected with the chytrid pathogen Bd in phonotaxis tests. Female túngara frogs prefer the calls of uninfected males (Bd-, open symbols) vs the calls of the same males when infected with Bd (Bd+, filled symbols). Proportion of female choice is shown for three types of calls: A) natural (n=67), B) amplitude normalized and amplitude inverted (n=87).

**Table 3.**
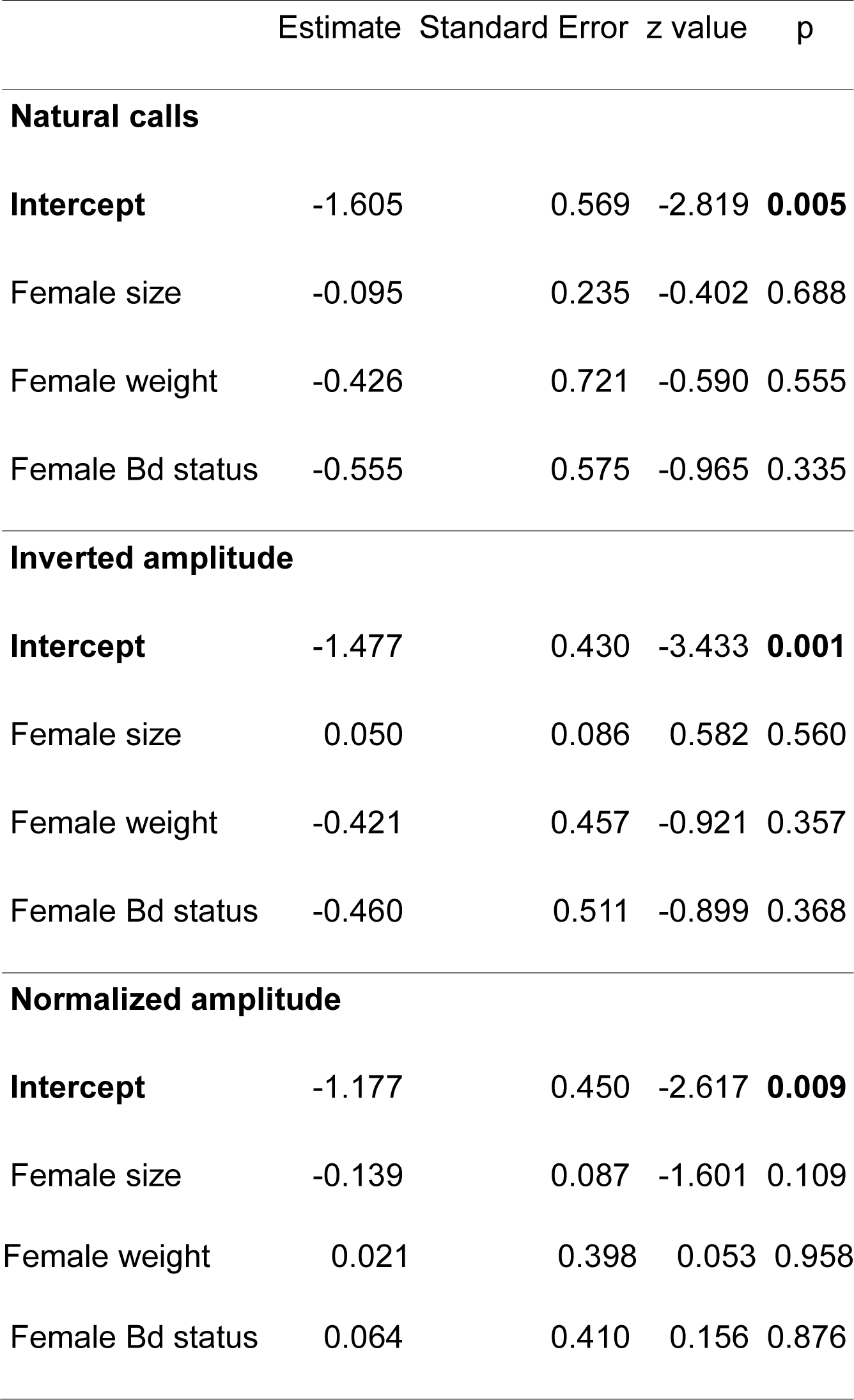
Summary of the general linear mixed-effects model to test for the effect of Bd infection status of males on female preference for the three experiments: natural calls, inverted amplitude, and normalized amplitude.

To determine if call amplitude was the salient feature influencing female mate choice, in the second and third experiments we manipulated the amplitude of the calls independent of other spectral or temporal characteristics of the acoustic signal. A total of 87 females were tested in these two experiments, and responded to all of the 6 tests for a total of 522 phonotaxis choices. When the calls were normalized, the females still preferred the calls of the males in their uninfected state (Fig. 3B). Similarly, when the call amplitudes were inverted, such that the call produced while the male was infected with Bd was of higher amplitude, the females still preferred the call produced after the infection was cleared (Fig. 3C).

## Discussion

High amplitude calls are attractive to females, however they are more energetically costly to produce (Ryan 1988), thus we expect males in poor condition to produce less attractive calls than healthier males in better condition. We predicted that infection with Bd would affect the males energetic state and therefore influence the amplitude of the males’ mating call. Females’ preference for the natural calls of an uninfected-male is not surprising since Bd-infected males called at lower amplitudes (Table 1) and female frogs often are attracted to calls with higher amplitudes (Gerhardt and Huber 2002). Consistent with the results of the tests with the natural calls, these preferences remained the same for each male even when we normalized or inverted the amplitude in the calls: females preferred the call of the male when it was clear of the infection over the call of the same male when infected (Table 3). These results show that, in addition to the amplitude of the mating call, other components of the call contribute to female mating preference for uninfected males.

Here we show that the effect on the properties of the males’ mating call of túngara frogs caused by the infection with Bd is salient to females and they discriminate against these calls. The female preference for calls of uninfected males suggests that there is an associated cost of infection with Bd in the male’s performance when producing a mating call. Thus infection with Bd incurs a cost in the attractiveness of mating calls of male túngara frogs, and might reduce their fitness. Since Bd infects amphibians’ skin, mating with an infected male increases the probability of infection. Females that discriminate against Bd-infected males may decrease their chance of sexual transmission of Bd. Other studies have suggested that infection with Bd changes calling behavior, but contrary to our results they suggest that the calls of males infected with Bd might be more attractive. The ultimate effect on female preference, however, has not been tested. In the Japanese tree frog, *Hyla japonica,* males infected with Bd produced more pulses per note and longer note duration suggesting a higher calling effort when infected (An and Waldman 2016). The Australian frog *Litoria rheocola*, is more likely to be found calling when males are infected and in good condition (Roznik et al. 2015), although the properties of the mating call did not differ between infected and uninfected males (Greenspan et al. 2016). Although both studies suggest that these changes in calling behavior due to infection with Bd may increase the males’ attractiveness, and therefore increase the dispersion of Bd, female preference tests were not performed so these assertions remain to be tested. Our study shows evidence that sub-lethal effects of infection with Bd on apparently resistant species should decrease males’ Darwinian fitness by making them less attractive to females, and may aid females in avoiding other potential sub-lethal effects of chytridiomycosis.

## Acknowledgments

Areli Benito and Jianguo Cui for assistance in field experiments, David Rodriguez for facilitating qPCR assays, and the STRI for logistical support.

## Data Accessibility Statement

Analyses reported in this article can be reproduced using the data provided by Rodríguez-Brenes et al. (2017).

## References

An D, Waldman B. 2016. Enhanced call effort in Japanese tree frogs infected by amphibian chytrid fungus. Biol. Lett. 12:20160018.

Bataille A, Cashins SD, Grogan L, Skerratt LF, Hunter D, McFadden M, Scheele B, Brannelly LA, Macris A, Harlow PS, Bell S, Berger L, Waldman B. 2015. Susceptibility of amphibians to chytridiomycosis is associated with MHC class II conformation. Proc. R. Soc. Lond. B Biol. Sci. 282:20143127.

Bates D, Mächler M, Bolker B, Walker S. 2015. Fitting linear mixed-effects models using lme4. J. Stat. Softw. 67:1–48.

Boyle D, Boyle D, Olsen V, Morgan J, Hyatt A. 2004. Rapid quantitative detection of chytridiomycosis (*Batrachochytrium dendrobatidis*) in amphibian samples using real-time Taqman PCR assay. Dis. Aquat. Organ. 60:141–148.

Brannelly LA. 2014. Reduced itraconazole concentration and durations are successful in treating *Batrachochytrium dendrobatidis* infection in amphibians. J. Vis. Exp. JoVE.

Ceballos G, Ehrlich PR, Barnosky AD, García A, Pringle RM, Palmer TM. 2015. Accelerated modern human–induced species losses: Entering the sixth mass extinction. Sci. Adv. 1:e1400253.

Chatfield MWH, Brannelly LA, Robak MJ, Freeborn L, Lailvaux SP, Richards-Zawacki CL. 2013. Fitness consequences of infection by *Batrachochytrium dendrobatidis* in northern leopard frogs (*Lithobates pipiens*). EcoHealth 10:90–98.

Collins JP, Crump ML. 2009. Extinction in Our Times: Global Amphibian Decline. Oxford University Press.

Ellison AR, Tunstall T, DiRenzo GV, Hughey MC, Rebollar EA, Belden LK, Harris RN, Ibáñez R, Lips KR, Zamudio KR. 2015. More than skin deep: functional genomic basis for resistance to amphibian chytridiomycosis. Genome Biol. Evol. 7:286–298.

Fernández-Beaskoetxea S, Bosch J, Bielby J. 2016. Infection and transmission heterogeneity of a multi-host pathogen (*Batrachochytrium dendrobatidis*) within an amphibian community. Dis. Aquat. Organ. 118:11–20.

Fites JS, Ramsey JP, Holden WM, Collier SP, Sutherland DM, Reinert LK, Gayek AS, Dermody TS, Aune TM, Oswald-Richter K, Rollins-Smith LA. 2013. The invasive chytrid fungus of amphibians paralyzes lymphocyte responses. Science 342:366–369.

Gerhardt HC, Huber F. 2002. Acoustic Communication in Insects and Anurans: Common Problems and Diverse Solutions. University of Chicago Press.

Greenspan SE, Roznik EA, Schwarzkopf L, Alford RA, Pike DA. 2016. Robust calling performance in frogs infected by a deadly fungal pathogen. Ecol. Evol. 6:5964–5972.

Guimarães M, Corrêa DT, Filho SS, Oliveira TAL, Doherty PF, Sawaya RJ. 2014. One step forward: contrasting the effects of toe clipping and PIT tagging on frog survival and recapture probability. Ecol. Evol. 4:1480–1490.

Harris RN, Brucker RM, Walke JB, Becker MH, Schwantes CR, Flaherty DC, Lam BA, Woodhams DC, Briggs CJ, Vredenburg VT, Minbiole KPC. 2009. Skin microbes on frogs prevent morbidity and mortality caused by a lethal skin fungus. ISME J. 3:818–824.

Hyatt A, Boyle D, Olsen V, Boyle D, Berger L, Obendorf D, Dalton A, Kriger K, Hero M, Hines H, Phillot R, Cambell R, Marantelli G, Gleason F, Colling A. 2007. Diagnostic assays and sampling protocols for the detection of *Batrachochytrium dendrobatidis*. Dis. Aquat. Organ. 73:175–192.

Kolbert E. 2014. The Sixth Extinction: An Unnatural History. Henry Holt and Company.

Kosch TA, Bataille A, Didinger C, Eimes JA, Rodríguez-Brenes S, Ryan MJ, Waldman B. 2016. Major histocompatibility complex selection dynamics in pathogen-infected túngara frog (*Physalaemus pustulosus*) populations. Biol. Lett. 12:20160345.

Kriger KM, Hero J, Ashton KJ. 2006. Cost efficiency in the detection of chytridiomycosis using PCR assay. Dis. Aquat. Organ. 71:149–154.

Madelaire CB, José da Silva R, Ribeiro Gomes F. 2013. Calling behavior and parasite intensity in treefrogs, *Hypsiboas prasinus*. J. Herpetol. 47:450–455.

R Core Team. 2015. R: A Language and Environment for Statistical Computing. Vienna, Austria: R Foundation for Statistical Computing.

Rachowicz LJ, Vredenburg VT. 2004. Transmission of *Batrachochytrium dendrobatidis* within and between amphibian life stages. Dis. Aquat. Organ. 61:75–83.

Rand AS, Ryan MJ. 1981. The adaptive significance of a complex vocal repertoire in a neotropical frog. Z. für Tierpsychol. 57:209–214.

Rodríguez-Brenes S, Rodriguez D, Ibáñez R, Ryan MJ. 2016. Spread of amphibian chytrid fungus across lowland populations of túngara frogs in Panamá. PLOS ONE 11:e0155745.

Roznik EA, Sapsford SJ, Pike DA, Schwarzkopf L, Alford RA. 2015. Condition-dependent reproductive effort in frogs infected by a widespread pathogen. Proc R Soc B 282:20150694.

Ryan MJ. 1985. The Túngara Frog: A Study in Sexual Selection and Communication. University of Chicago Press.

Ryan MJ. 1988. Constraints and patterns in the evolution of anuran acoustic communication. In: Fritzsch, B., Ryan, M., Wilczynski, W., Walkowiak, W., Hetherington, T., editors. The Evolution of the Amphibian Auditory System. New York: John Wiley and Sons Inc. p. 637– 677.

Ryan MJ, Rand AS. 1990. The sensory basis of sexual selection for complex calls in the Túngara frog, *Physalaemus pustulosus* (sexual selection for sensory exploitation). Evolution 44:305–314.

Savage AE, Zamudio KR. 2011. MHC genotypes associate with resistance to a frog-killing fungus. Proc. Natl. Acad. Sci. 108:16705–16710.

Venesky M, Parris M, Storfer A. 2009. Impacts of *Batrachochytrium dendrobatidis* infection on tadpole foraging performance. EcoHealth 6:565–575.

Voyles J, Young S, Berger L, Campbell C, Voyles WF, Dinudom A, Cook D, Webb R, Alford RA, Skerratt LF, Speare R. 2009. Pathogenesis of chytridiomycosis, a cause of catastrophic amphibian declines. Science 326:582–585.

Wake DB, Vredenburg VT. 2008. Colloquium Paper: Are we in the midst of the sixth mass extinction? A view from the world of amphibians. Proc. Natl. Acad. Sci. 105:11466–11473.

